# Differential expansion microscopy

**DOI:** 10.1101/699579

**Authors:** Sebastian P. Pernal, Asiri Liyanaarachchi, Domenico L. Gatti, Brent Formosa, Rishika Pulvender, Eric R. Kuhn, Rafael Ramos, Akshata R. Naik, Kathleen George, Suzan Arslanturk, Douglas J. Taatjes, Bhanu P. Jena

## Abstract

Expansion microscopy (ExM) involves the use of hydration-competent polymers to physically expand biological specimens approximately 4-fold linear increase to achieve 70 nanometer resolution using an ordinary diffraction limited optical microscope. Optimal conditions however for antigen retention during the expansion process and the relative expansion between organelles within cells has remained unclear. It is reported that different tissues expand to different extents, suggesting that although isotropic expansion is believed to occur, different subcellular compartments with different composition would undergo anisotropic or differential expansion (DiEx). Consequently, there would be distortion of the native shape and size of subcellular compartments upon expansion, parameters which are critical in assessing cellular states in health and disease. Here we report optimal fixation and expansion conditions that retain structural integrity of cells while exhibiting up to 8-fold linear and therefore 512-fold volumetric expansion. Anisotropic expansion is observed not just between tissues, but between different subcellular compartments and even within subcellular compartments themselves. Combining image analysis and machine learning, we provide an approach for the rapid and precise measurement of cellular and subcellular structures in expanded tissue. Using both manual and computation assessment of morphometric parameters, we demonstrate expansion to be anisotropic and therefore refer to this this method as differential expansion microscopy (DiExM).

## Introduction

An average human cell is composed of nearly a billion protein molecules, and millions of lipids, carbohydrate and nucleic acids, present in a wide array of combinatorial patterns to establish intracellular organelles and nanomachines that perform various cellular tasks. Elucidating the cellular organization of these biomolecules is key to understanding life processes and in the detection and treatment of disease. Since the invention of the light microscope over 400 years ago, a wide range of imaging tools have been invented and utilized to study the cell. In the past 75 years, transmission electron microscopy (TEM) and the atomic force microscope (AFM) have played a pivotal role in the discovery of cellular organelles^1^ and nanomachines such as ribosome^2^, and the porosome^3-6^, as well as and elucidation of the interactions between membrane-associated fusion proteins called SNAREs to establish continuity between opposing lipid membranes^7,8^. Two further powerful tools have recently emerged, namely cryo-electron microscopy (Cryo-EM)^9^ and super-resolution fluorescence microscopy^10,11^, both capable of resolving single molecules. In contrast to the development of such high-resolution imaging systems, recently for the first time, the substrate was enlarged to enable nanoscale imaging using an ordinary diffraction limited light microscope^12^. In this simple yet novel new approach termed expansion microscopy (ExM)^12^, hydration-competent polymers are used to physically expand biological specimens allowing imaging at approximately 70 nanometer resolution using a light microscope^12-14^. Earlier studies^14^ reported that different tissues expand to different extents, for example, an expansion factor of 4.6 was observed for human lung tissue compared to 4 for human breast tissue under identical expansion conditions^14^, suggesting that although isotropic expansion is believed to occur, different subcellular compartments with different composition would undergo anisotropic or differential expansion (DiEx). Consequently, there would be distortion of the native shape and size of subcellular compartments upon expansion, parameters which are critical in assessing cellular states. Furthermore, while the current protocol^12^ provides a 4-fold linear expansion translating to a 64-fold volumetric increase, this limit had yet to be overcome while retaining the structural integrity of cells. Additionally, the optimal conditions for antigen retention during the expansion process and the relative expansion between organelles within cells had not been explored.

## Results and Discussion

Here we report a further developed version of ExM which is (i) demonstrating greater than 500-fold volumetric expansion (Fig. 1), (ii) is anisotropic between tissues, organelles and within organelles, and (iii) exploits machine learning to assess the resulting differential expansion. We term this approach, differential expansion microscopy (DiExM). A major concern in ExM is the loss of cellular antigens during the expansion process. Prefixation of tissues and cells therefore using 4% paraformaldehyde (PFA) have previously been adopted in ExM studies^12-14^. While increasing the concentration of fixative would help retain cellular integrity and minimize loss of cellular antigens during the ExM process, it would result in decreased expansion due to more crosslinking between cellular antigens. A balance between the extent of fixation and expansion is therefore necessary to obtain meaningful intracellular information from expanded tissue. Similarly, to achieve optimal expansion while retaining structural integrity would necessitate the uniform introduction of the gel-forming monomer into cells prior to polymerization and formation of a polyelectrolyte gel. Furthermore, a balance needed to be reached, where optimal tissue digestion would enable maximal expansion while retaining cellular antigen content and structural integrity. To achieve these objectives, rat liver tissue was prefixed using increasing concentration of PFA (1%, 2%, 4% and 8%) followed by a modified ExM approach implementing optimal monomer loading of the tissue, followed by polymerization, limited digestion, and expansion. As predicted, increased concentration of PFA resulted in retention of the structural integrity of cells following expansion, as observed using transmission electron microscopy (TEM) (Fig. 1a). A 1% PFA fixation resulted in more loss of cytoplasmic content following expansion, suggesting a higher concentration of PFA might be optimal. Although exposure to increased concentration of PFA might in principle resulted in decreased expansion, fixation using 4% PFA demonstrated maximum expansion of up to 8-fold linearly and 512-fold volumetrically (Fig. 1). Since increased concentration of PFA beyond 2% resulted in no detectable changes to cellular morphology assessed by TEM, the 4% PFA fixation protocol was adopted (Fig. 1a). Analysis of the distributions of the linear fold-expansion of hepatocyte nucleus and cell body demonstrate a 2-to 8-fold expansion, with the 4% PFA-fixed tissue demonstrating maximal expansion. While the mean linear expansion factor of the hepatocyte nucleus ranges from 2.16 to 3.86, the mean linear expansion factor for the entire hepatocyte, ranges from 2.49-to 4.71, demonstrating anisotropic expansion between cellular compartments (Fig. 1b, c). These results are not unexpected since different cell types having different composition also demonstrate anisotropic or differential expansion as mentioned earlier (expansion factor of 4.6 for human lung tissue compared to 4 for human breast tissue)^14^.

**Figure 1.**
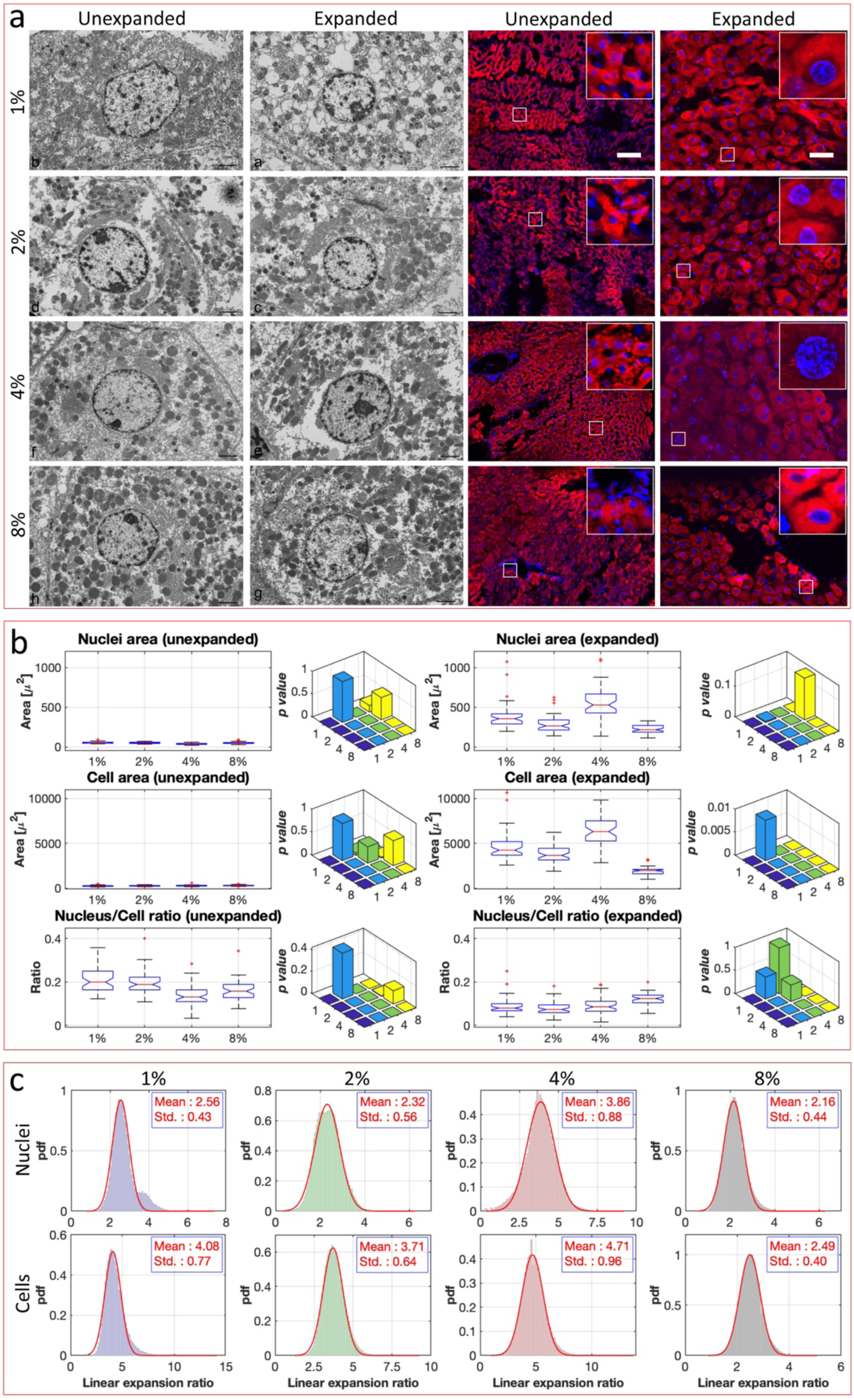
Modified approach using sodium polyacrylate-induced up to 8-fold linear expansion of rat hepatocytes prefixed in 4% para formaldehyde (PFA), corresponding to a 512-fold increase in cell volume. **a**, (Left two panels) Representative transmission electron micrographs of the unexpanded and expanded rat liver tissue demonstrating no visible change between the 2%, 4% and 8% PFA fixed tissue, while the 1% expanded cell reflects some loss of cytosolic material. *(Scale bar = 2 µm)*. (Right two panels) Light fluorescent micrographs of rat liver sections pre-fixed in different concentrations of PFA and stained using R18 dye for cellular lipids and DAPI blue for the nucleus, demonstrates maximum expansion of up to 8-fold in linear expansion in the 4% PFA-fixed hepatocytes. *(Scale bar = 100 µm)*; randomly chosen areas (boxed) from within each of the micrographs are magnified 5-fold and placed as insets on the top right corner of each of the fluorescent images. Note maximal expansion of both the cell and the nucleus in hepatocytes fixed using 4% PFA. **b**, Significant changes in the cell and nuclear area and the ratios between them in unexpanded and expanded cells are shown as box plots. Box plots representing one-way analysis of variance (ANOVA) between the feature distributions of nuclear areas, cellular areas, and nucleus/cell area ratios in unexpanded and expanded cells that were hand selected for having the cell contour clearly demarcated from surrounding cells. In each box, the central mark is the median and the edges of the box are the 25th and 75th percentiles. The whiskers extend to the most extreme data points that are not considered outliers. The outliers are plotted individually as red + signs outside the whiskers. The interval endpoints are the extremes of the notches. The extremes correspond to *q*2 – 1.57(*q*3 – *q*1)/sqrt(*n*) and *q*2 + 1.57(*q*3 – *q*1)/sqrt(*n*), where *q*2 is the median (50th percentile), *q*1 and *q*3 are the 25th and 75th percentiles, respectively, and *n* is the number of observations. Two medians are significantly different at the 5% significance level if their intervals do not overlap. Large differences in the center lines of the boxes correspond to large *F*-statistic values and correspondingly small *p*-values. Pairwise *p-values* for the four different concentrations of fixative tested (1, 2, 4, 8%) are shown as a 3D-bar plot on the right side of each set of box plots for a particular feature considered. Note, the maximum expansion in the 4% PFA-fixed tissue. **c**, Distribution of the linear fold expansion in the hepatocyte nucleus and cell body demonstrates from 2-to 8-fold expansion, with the 4% PFA-fixed cell demonstrating maximal expansion. Histograms are normalized to represent a probability density function (pdf) with unit area. A fit of a normal distribution with given mean and standard deviation (as shown in the inset) is displayed as a red line superimposed on each histogram. Note the average expansion factor of the nucleus compared to the cell is different for every percentage of fixative used.

To further determine if expansion is anisotropic, one of the most fibrous among cells, the human skeletal muscle cell, was expanded next. Similar to the differential expansion observed between human lung and breast tissue^14^, human skeletal muscle cells demonstrate anisotropic expansion (Fig. 2) compared to rat hepatocytes (Fig. 1). Anisotropic expansion is also observed in the nucleus and myosin fibers of the human skeletal muscle cell (Fig. 2). While a 3.5-fold average linear increase in the size of the nucleus is observed in the expanded compared to unexpanded cells, there is a modest 2.7-fold increase in the average myosin fiber width following expansion, demonstrating anisotropic expansion between organelles in human skeletal muscle cells. To assess whether the 3.5-fold linear expansion of the nucleus is uniform in every direction (i.e., isotropic), the ratio of the nuclear length to width was estimated in the expanded nuclei and compared to unexpanded. Ratio of nucleus length to width in the unexpanded cells was found to be 1.82 (1.82 ± 0.05; n=50) and in the expanded 2.33 (2.33 ± 0.09; n=50), a significant (p<0.0001) 28% increase in the expanded cells, demonstrating anisotropic expansion of the nucleus (Fig. 2k). To assess if the longitudinal expansion of the nucleus is a consequence of the variations in chemical composition at different domains within the nucleus, bright areas of nuclear staining with 4’,6-diamidino-2-phenylindole (DAPI) were assessed. DAPI blue-fluorescent DNA stain binds with high specificity to AT regions of double stranded DNA (dsDNA) and is commonly used to label the cell nucleus^15^. DAPI binds to intranuclear heterochromatin regions^15^ but not to the nucleolus, the site for ribosome biogenesis. Intense DAPI staining from non-intensely stained regions within the nucleus is therefore a reflection of differences in intranuclear composition. We utilized this differential staining of the nucleus by DAPI to investigate whether different regions of the nucleus expand differently. The ratio of the nuclear area intensely labeled with DAPI to the entire nuclear area decreases following expansion (Fig. 2l). In the unexpanded cells the ratio of DAPI bright regions to the entire nuclear area was found to be 0.3 (0.3 ± 0.017; n=50), as opposed to 0.224 in the expanded cells (0.224 ± 0.017; n=50), showing a significant (p<0.0001) 25% decrease in the expanded cells. These results demonstrate that intranuclear domains have different composition, and therefore expand differently. The nucleus therefore expands in an anisotropic manner (Fig. 2k, 2l). Areas within the nucleus of human skeletal muscle cells with high concentration of DAPI-binding dsDNA, expand relatively less than the rest of the nucleus and therefore demonstrate differential expansion (Fig. 2k, 2l). Additionally, this new expansion protocol demonstrates retention of the immuno-fluorescently labeled integrity of cells.

**Figure 2.**
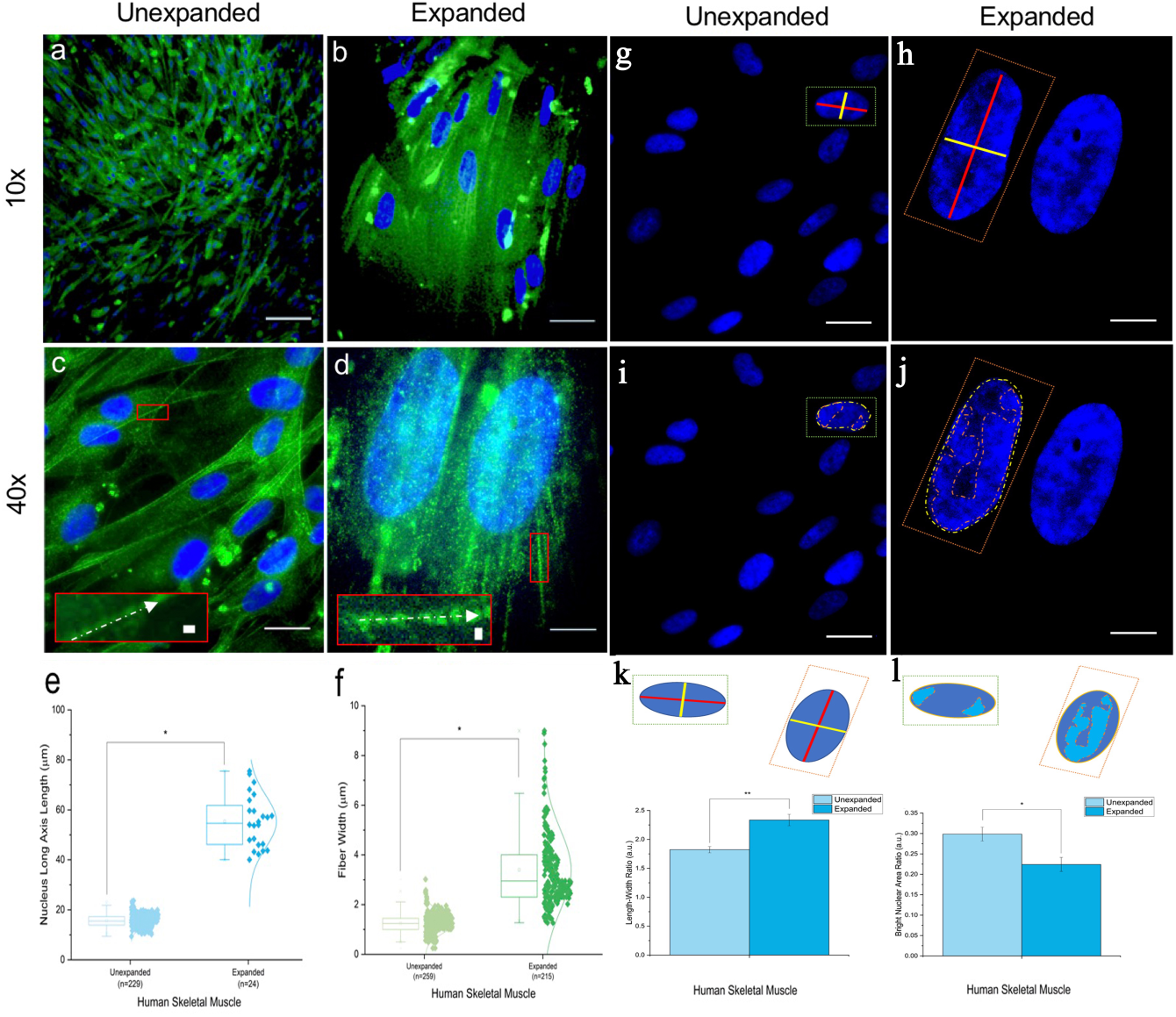
Human skeletal muscle cells undergo differential expansion. **a-f**, Differential expansion of the nucleus compared to myosin IIb-specific fiber thickness. Staining of the nucleus with DAPI and immunolabeling of myosin IIb in human skeletal muscle cells followed by expansion, demonstrates a 3.5-fold linear increase in the size of the nucleus as opposed to a 2.7-fold increase in myosin fiber width. **a**,**c**, Unexpanded and **b**,**d**, expanded human skeletal muscle cells labeled with anti-myosin IIb (green) and DAPI for nuclei (blue), scale bar 20µm. **e**,**f**, Size of the nucleus in unexpanded (15.71 ± 2.56µm; n=229) and expanded cells (55.36 ± 10.66µm; n=24) demonstrates a significant 3.5-fold increase (p < 0.0001). **f**, Similarly, the size of myosin fiber thickness in unexpanded (1.24 ± 0.42 µm; n=259; inset 1µm) and expanded muscle cells (3.40 ± 1.62 µm; n=215; inset 1µm) demonstrates a significant 2.7-fold increase (p < 0.0001). Note, while there is a 3.5-fold linear increase in the size of the nucleus, only a 2.7-fold increase in myosin fiber width is observed in expanded human skeletal muscle cells, demonstrating differential expansion of the two organelles. **g-n**, Anisotropic expansion of the human skeletal muscle cell nucleus. **g**, High magnification unexpanded and **h**, high magnification expanded image of human skeletal muscle cells labeled with DAPI. **i**,**j**, Ratio of nucleus length to breadth in the unexpanded cells is (1.82 ± 0.05; n=50) and in the expanded (2.33 ± 0.09; n=50), demonstrating a significant 28% increase (p < 0.0001) in the expanded cells, hence the anisotropic expansion of the nucleus. **k-n**, In contrast, the ratio of the nuclear area intensely labeled with DAPI compared to the entire nucleus area decreases following expansion. In the unexpanded cells the ratio of DAPI bright regions to the entire nucleus area is (0.3 ± 0.017; n=50) and in the expanded (0.224 ± 0.017; n=50), demonstrating a significant 25% decrease (p < 0.0001) in the expanded cells, further demonstrating that intranuclear domains with different chemistry, i.e., areas with high concentration of DAPI-binding dsDNA, expand relatively less than the rest of the nucleus.

Changes in the ratio of size and shape between subcellular organelles and the concentration and distribution of various cellular antigens are key parameters that reflect the health status of cells. A major issue therefore is the anisotropic expansion of intracellular compartments. To account for the anisotropic expansion and to recover the high-resolution and proportional images of expanded cells, we employed machine learning. To this end, we developed a Matlab^®^ based interactive application for the analysis of control and DiExM images of human skeletal muscle cells and rat liver cells (Fig. 3, Fig. S1-3 in Supplementary Information and video/audio tutorial). The application executes sequentially different layers of image processing and analysis. First, it identifies each organelle (through segmentation) and then extracts quantitative information about various features (i.e., feature engineering) of nuclei, cytoplasm, and other imaged molecules or organelles of importance. This machine learned morphometric analysis has confirmed the anisotropic expansion of the human skeletal muscle and rat liver cells. In light of the different levels of expansion undergone by different cellular compartments, this quantitative information is extremely valuable to interpret the DiExM images of cells properly in the study of cellular structure-function and in pathologic states that affect cellular and sub-cellular dimensions. We anticipate that different diseased states of human cells, with their effects on the biochemical and structural properties of cellular components, will be reflected in different levels of anisotropic expansion. As a proof of concept of the capacity of a machine learning approach to recognize differential changes in expansion, and thus possible different diseased states, we have trained an artificial neural network (NN) using images of both unexpanded and expanded liver cells prefixed in either 1, 2, 4, and 8% PFA. Upon training, the NN was capable of classifying unlabeled images as belonging to one of these five classes with 84% accuracy (Supplementary Figs. S2, S3). This clearly shows that the quantitative information extracted from the features of the cells is capable of distinguishing different levels of expansion. We anticipate that these features can further be used to differentiate the health status of cells as diseased states may result in different levels of expansion.

**Figure 3.**
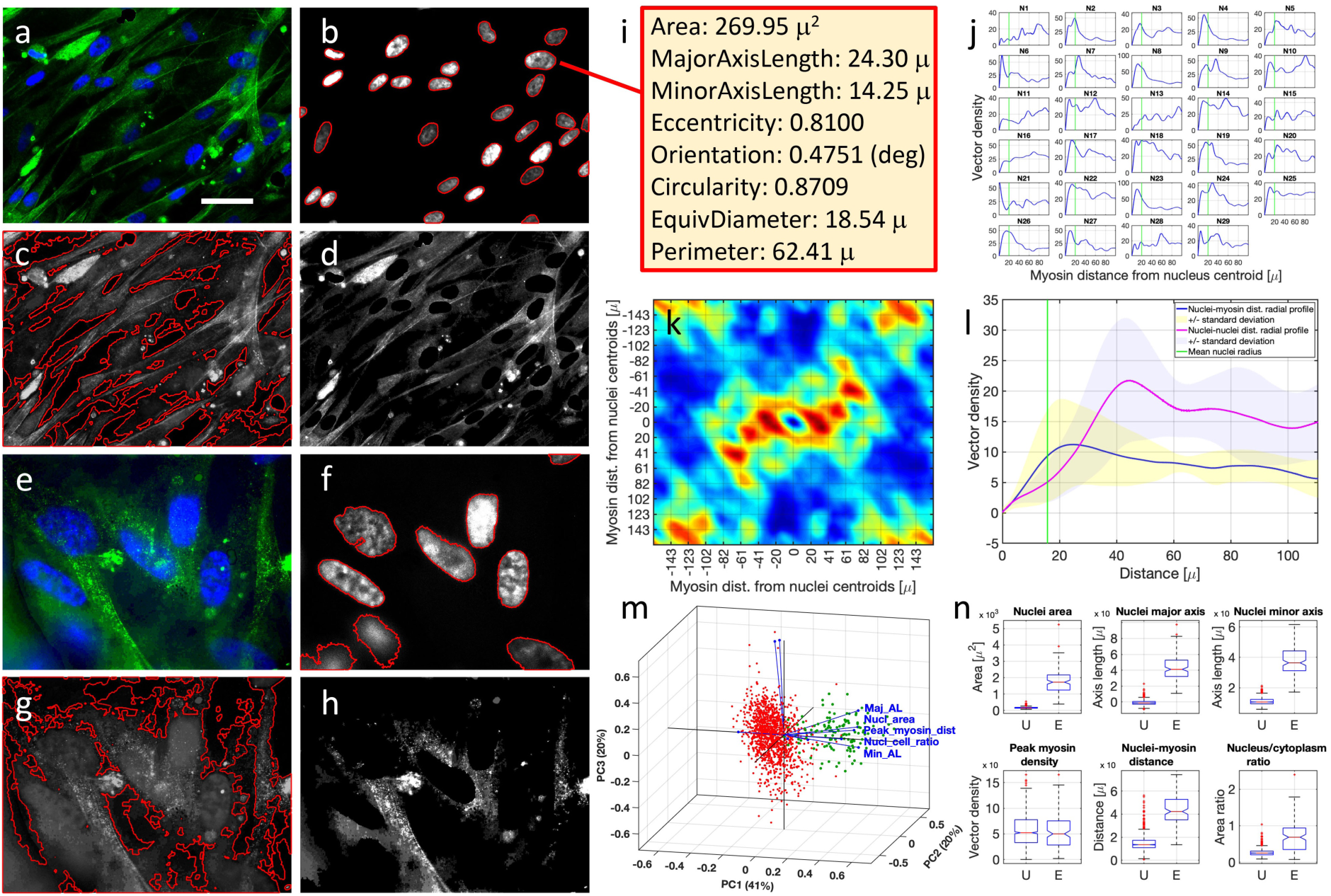
Machine learned morphometric parameters confirm anisotropic expansion of the human skeletal muscle cell. **a,e**, Unexpanded control cells and expanded cells (respectively) labeled with anti-myosin IIb (green) and DAPI for nuclei (blue); scale bar 40µm. **b,f**, DAPI channel of the same images in panels **a** and **e**, respectively, with machine learning derived nuclei contours highlighted in red. **c,g**, Gray scale representation of the same images of panels a and e, respectively, with machine learning derived cell contours highlighted in red. **d,h**, Anti-myosin IIb channel of the same images in panels **a** and **e**, respectively. **i**: Some machine learned features of the nucleus in panel **b** pointed by the red line. Circularity specifies the roundness of objects and is computed as (4*Area*pi)/(Perimeter^2^). For a perfect circle, the circularity value is 1. Given an elliptical nucleus, its eccentricity is the ratio of the distance between the foci of the ellipse and its major axis length. An ellipse whose eccentricity is 0 is actually a circle, while an ellipse whose eccentricity is 1 is a line segment. **j**: Each inset shows the profile of the intensity (blue trace) of the myosin signal with respect to the centroid of each nucleus (labeled as N1, N2, …,N29) of the image shown in panel **a**, as calculated from a radially averaged Difference Patterson map (Patterson map of (nucleus + myosin) – (Patterson map of nucleus + Patterson map of myosin)) (see Methods). A vertical green line in each panel marks the mean radius of all nuclei in the image. The mean density distribution of the myosin with respect to all the nuclei centroids is calculated by averaging the individual profiles for each nucleus. **k**, Difference Patterson map (Patterson map of (all nuclei + myosin) – (Patterson map of all nuclei + Patterson map of myosin)) of the image shown in panel **e**. The center of the image is the center of all nuclei. Areas of the map with dark blue and dark red color reflect low and high vector densities, respectively. Regions of the map with high vector density reflect a strong myosin signal. Radial average of this map provides the distribution of myosin around the nuclei centroids. **l**, Mean density distribution of myosin with respect to all the nuclei centroids (blue trace with ±1 standard deviation boundary as a shaded yellow area), and of each nucleus centroid with respect to all the nuclei centroids (magenta trace with ±1 standard deviation boundary as a shaded lilac area) in the image shown in panel **e**, as calculated by averaging the individual profiles for each nucleus. A vertical green line marks the mean radius of all nuclei in the image. The myosin peak is located at approximately 24 µm from the nuclei centroids (blue trace), while the mean nucleus to neighbor nucleus distance is approximately 43 µm. **m**, Principal Component Analysis of machine learned nuclear features in unexpanded (red circles) and expanded (green circles) cells. Nuclei features included in the analysis were: area, eccentricity, major axis length, minor axis length, axes length ratio, mean distance from all other nuclei, mean distance from neighbor nuclei, myosin peak distance, myosin peak density, nucleus/cell area ratio. The first 3 principal components account for 81% of the data variance. Discrimination between unexpanded and expanded cells occurs primarily along the 1^st^ PC axis. Blue vectors in PC space represent the directions along which different real space features are most discriminating: among these, nuclear axes length, nuclear area, nuclear/cell area ratio, and peak myosin distance from nuclei centroids are the features that align best with the 1^st^ PC axis. **n**, Box plots representing one-way analysis of variance (ANOVA) between the distributions of machine learning acquired nuclei features in unexpanded (U label on the x-axis) and expanded (E label on the x-axis) cells. In each box, the central mark is the median and the edges of the box are the 25th and 75th percentiles (1st and 3rd quantiles). The whiskers extend to the most extreme data points that are not considered outliers. The outliers are plotted individually as red + signs outside the whiskers. The interval endpoints are the extremes of the notches. Two medians are significantly different at the 5% significance level if their intervals do not overlap. Large differences in the center lines of the boxes correspond to large *F*-statistics and correspondingly small *p*-values. *p-values* for features shown in panel **n** were all ≪ 10^−10^ with the exception of the myosin peak density (*p* ≅ 0.5), whose distributions in unexpanded and expanded cells almost completely overlap.

## Summary

In summary, we have developed a modified expansion protocol (Fig. 4, Supplementary material) to obtain up to >500-fold volumetric expansion of biological tissues, while retaining the structure and the immuno-fluorescently labeled integrity of cells. We demonstrate that expansion is anisotropic, exhibiting differential expansion between tissues, cellular organelles and even within within organelles themselves. Both manual and machine learned morphometric analyses confirm anisotropic expansion of the human skeletal muscle and rat liver cells examined in the current study. Finally, we show that an artificial NN based analysis using DiExM images is successful in recognizing features of differential expansion that may be characteristic of particular diseased states.

**Figure 4.**
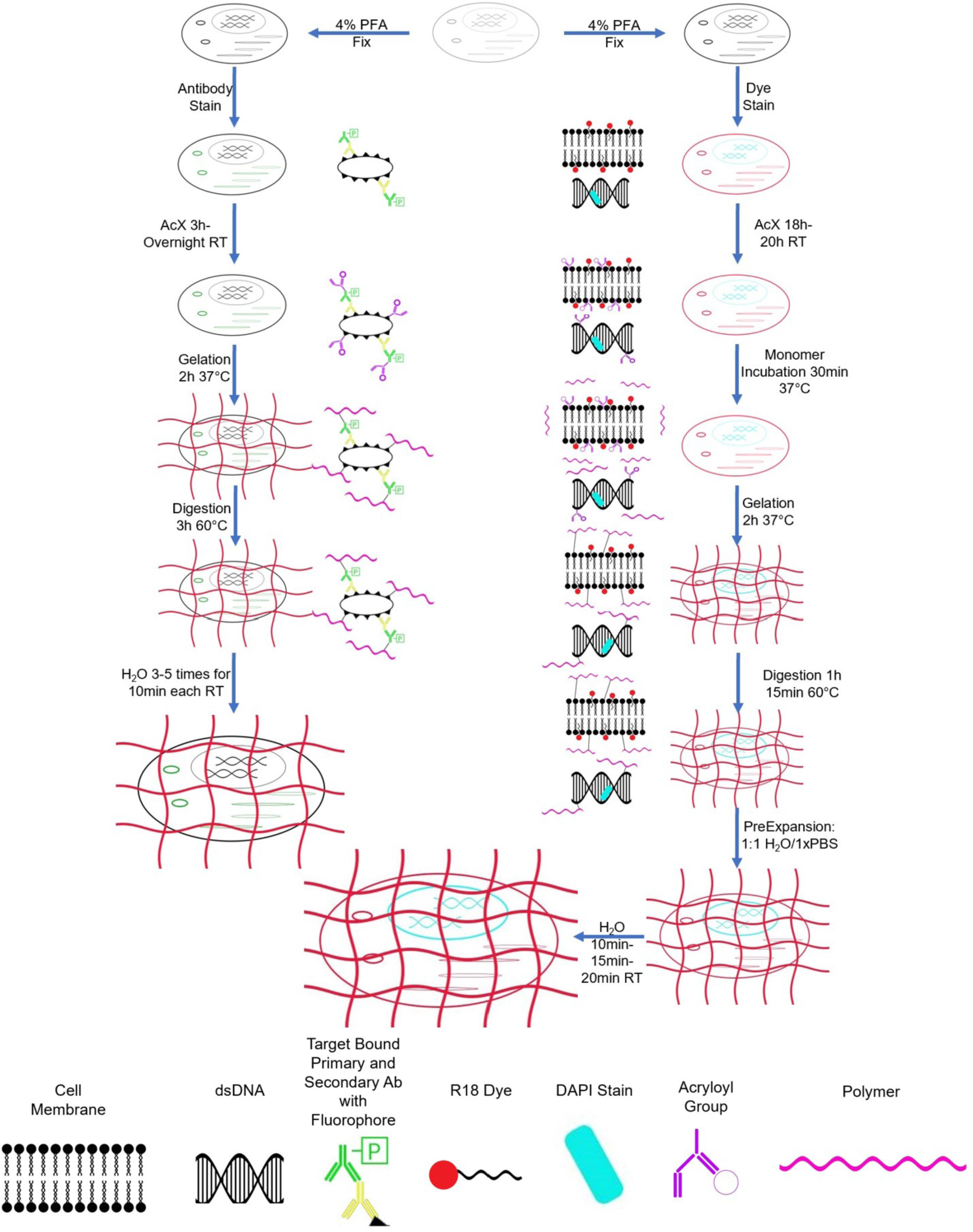
Flow chart depicting previously published protocol and the new protocol used for this paper. *Left*, Y. Zhao *et al.* (2017) expansion protocol^14^; Samples chemically fixed using 4% paraformaldehyde (PFA), stained with antibodies using standard immunostaining protocols, 6h-overnight AcX treatment to anchor, 2h gelation at 37°C in 100% humidity, followed by 3h digestion at 60°C and subsequent expansion. *Right*, Samples fixed, stained with R18 dye and DAPI, 18-20h Acx treatment to anchor, 30 min monomer incubation prior to 2h gelation at 37°C in 100% humidity, followed by 1h 15 min digestion at 60°C, followed by PreExpansion overnight in 50% phosphate buffered saline (PBS) pH 7.4, followed by incubation in double distilled water for 10, 15 and 20 min each between changes of water.

## Supplementary Information

contains methods and three figures (1S-3S), link to a video/audio tutorial and associated references.

## Author Contributions

B.P.J. developed the idea, B.P.J., S.P.P., A.L. and D.L.G., designed experiments for the study. B.P.J. and D.L.G. wrote the manuscript. S.P.P., A.L., B.F., E.R.K., R.R. and K.G. performed the expansion studies. A.R.N. performed the human primary skeletal muscle cell cultures. D.L.G., S.A. and B.P.J. participated in the machine learning (ML) aspects of the study and D.L.G. performed all ML studies. D.J.T. performed electron microscopy. R.P. helped S.P.P., A.L., B.F., in expansion experiments and in the manual morphometric analysis of the images. All authors participated in discussions and proofreading the manuscript.

## Acknowledgements

Work presented in this article was supported in part by the National Science Foundation grants EB00303, CBET1066661 (BPJ).

