## Supplementary Material for "Differential expansion microscopy"

[1]*Department of Physiology,* [2]*NanoBioScience Institute,* [3]*Center for Molecular Medicine & Genetics,* [4]*Biochemistry, Microbiology and Immunology, School of Medicine;* [5]*Department of Computer Science, College of Engineering, Wayne State University, Detroit, MI 48201, USA* [6]*Department of Pathology and Laboratory Medicine, Microscopy Imaging Center, University of Vermont College of Medicine, Burlington, VT 05405, USA*

[†]These authors contributed equally to the study.

### Methods

#### 1. Differential Immuno Expansion Microscopy Protocol

**Tissue Processing**

Rat Liver tissue was isolated. Liver tissue were cut to uniform size and each fixed in PFA (either 1%,2%,4%, or 8%) for 24hrs at 4°C. Tissues was then washed with 1% PBS 3 times for 30mins at RT. Tissue set in OCT block and frozen at -80°C. When needed, 30μm sections were taken.

**Immuno Labeling & Staining**

Sections were blocked with MaxBlock blocking medium (Active Motif) containing 0.1% Tween-20 for 2 hours at RT, followed by 2 washes with 1X PBS for 5mins each at RT. Tissue sections were incubated with the appropriate dilution for each primary and fluorescent secondary antibody which was optimized for each tissue separately. Lipids in the tissue sections were labeled with Octadecyl Rhodamine B Chloride (R18) dye in 1X PBS (1μM), followed again by 2 washes with 1X PBS for 5mins each at RT. Sections were incubated with 4′,6-diamidino-2-phenylindole (DAPI) in 1X PBS (0.2μM), wash 2 times with 1X PBS containing 0.1% Tween-20 for 5mins each at RT.

**Anchoring Treatment**

Dissolve Acryloyl-X, SE (Acx, Thermo Fisher Scientific) in anhydrous DMSO to result in a 10mg/ml stock solution. Aliquot the stock solution in 20ul aliquots and store in a desiccated environment at -20C. Make a AcX solution with 1x PBS at a concentration of 0.1mg/ml and incubate for 18-20 hours at room temperature.

**Polymer Synthesis**

Stock solutions for each component of the monomer solution was prepared and stored. A monomer solution made Sodium acrylate at 8.6% (stock: 38% dissolved in 1x PBS stored at -20C, Sigma), acrylamide at 2.5% (stock: 50% dissolved in 1x PBS stored at

4°C), N,N'-Methylenebisacrylamid at .10% (stock: 2% dissolved in 1x PBS stored at 4°C, Sigma), Sodium chloride 11.7% (stock: 29.2% dissolved in 1x PBS stored at 4°C), 1x PBS (stock: 10x stored at Room temperature), and Water was prepared fresh each time from stock solutions. Sections were incubated with the monomer solution for 30 minutes at 37°C in humid conditions. Monomer solution was then removed, and fresh gelling solution was added to the samples. Gelling solution contains the monomer solution and the chemicals .5% 4-hydrozy-2,2,6,6-tetramehtylpiperidin-1-oxy (4HT, Sigma) as an inhibitor, .2% tetramethylethylenediamine (TEMED, Sigma) as an accelerator, and .2% ammonium persulfate (APS, Sigma) as an initiator. 4HT, TEMED, and APS were added sequentially in that order to the gelling solution and samples were then immediately placed in a humidified water bath at 37°C for 2 hours.

**Sample Digestion**

After Gelation samples were incubated with 8 U/ml proteinase K (New England Biolabs) in a Digestion buffer consisting of 50mM Tris (pH8), 25 mM EDTA, .5% triton X-100, and 0.8M NaCl for 1 hour and 15 minutes at 60°C. after samples were covered with a 1:1 solution of 1x PBS and Deionized water for 12-15 hours.

**Expansion**

Excess solution was removed from the samples and do they could reach room temperature by leaving them out at room temperature for around an hour. Samples were washed 3 times with deionized water and incubated for 10, 15 and 20 minutes respectively.

**Imaging**

Sections were imaged on a Zeiss apotome imager Z1 fluorescence microscope at 10X and 40X magnifications. For each treatment group, 4 spatially distinct and randomly chosen 2-channel (DAPI 461nm and DsRed 568nm) images were acquired. Each image was extracted with Zeiss apotome imager with extended depth of focus rendered through Zen Imaging Software (Carl Zeiss AG).

### 2. Electron Microscopy

Transmission electron microscopy of unexpanded and expanded rat liver tissue prefixed for 24h in 1, 2, 4 and 8% PFA and then stored in 2% PFA until further processing was performed. Briefly, tissue pieces were further fixed in half-strength Karnovsky's fixative (1.5% glutaraldehyde, 1.0 % formaldehyde in 0.1M cacodylate buffer , for 80 minutes at 4°C, followed by post-fixation for 1h at 4°C in 1% $OsO_4$ in 0.1 M cacodylate buffer. The sample was then dehydrated in a graded series of ethanols, through propylene oxide and infiltrated and embedded in Spurr's resin. Ultrathin sections were cut with a diamond knife, retrieved ontogrids, and contrasted with alcoholic uranyl acetate and lead citrate. Grids were viewed with a JEOL 1400 transmission electron microscope (JEOL USA, Inc., Peabody, MA) operating at 80 kV, and digital images were acquired with an AMT-XR611 11 megapixel ccd camera (Advanced Microscopy Techniques, Danvers, MA).

### 3. Image Processing and Acquisition of Cellular Features

Image processing and acquisition of relevant features of unexpanded and DeXM expanded cell were carried out using an interactive application running in the Matlab environment and requiring the Image Analysis and Deep Learning Toolboxes. The complete Matlab code, test data, and a video tutorial on the use of the application are included as Supplementary material. The application executes sequentially different layers of image processing and analysis. Cropping, removal of local artifacts via a hand-drawn mask, and channel selection are user directed. Non-uniform background illumination is corrected by morphological opening [1]. The opening operation removes from the background all objects that cannot completely contain a disk-shaped structuring element with a user defined radius (typically the average nuclei radius): this background is then removed from the image. Uncorrelated channel color features (i.e, the blue of DAPI stained nuclei, the green of myosin decoration) are enhanced by decorrelation stretching (a technique affine to principal component analysis) based on the eigen decomposition of the channel-to-channel correlation matrix [2]: a stretch factor for each channel is derived as the inverse square root of the corresponding eigenvalue. Additional linear contrast stretching further expands the color range of the resulting image by saturating equal fractions at high and low intensities: typically, the transformed color range within each band is mapped to a normalized interval between 0.01 and 0.99, saturating 2%. Denoising is alternatively carried out by 1. bilateral filtering (very fast), 2. passing each channel of an RGB image (or the single luminance channel of an equivalent image in CIE 1976 Lab color space) through a denoising convolutional neural network (DnCNN) [3] pre-trained on images with added Gaussian noise (slow), or 3. Passing an RGB image through a pre-trained multiscale context aggregation convolutional neural network (CA_CNN) that is used to approximate an image filtering pipeline (intermediate in speed and outcome between 1 and 2) [4].

Nuclei segmentation was carried out on the B channel binarized by the Otsu method [5] with a combination of global and adaptive thresholding [6] with user defined sensitivity, followed by area opening (which removes all connected components with fewer than $p$ pixels from the binary image), cycles of dilation, hole filling and erosion of the nuclei mask. Mask pixels were defined as connected if their edges or corners touched. Two adjoining pixels were labeled as part of the same object if 8-connected, that is if they were both on and connected along the horizontal, vertical, or diagonal direction. In the case of partially overlapping nuclei or nuclei of difficult shape, visual inspection of the B channel was necessary to complement the nuclei masks with user defined separating segments and/or hand drawn polygonal mask elements (as shown in the Video Tutorial). Upon segmentation, morphological features extracted from individual nuclei included area, convex area, bounding box, centroid, pixel values with maximal, minimal, mean and s.d. intensity, weighted centroid based on pixels intensity, circularity, eccentricity, equivalent diameter, major and minor axis length, orientation, perimeter, solidity. The distribution of these parameters was further used to identify outliers likely to be artifacts or partial nuclei at the edge of an image, with the option to remove them, followed by mask updating.

Cell outlines were identified by a global/local-adaptive thresholding procedure similar to the one described for nuclei but carried out on a gray-scale image derived by combining the G channel (myosin) and B channel (nuclei) of the original RGB image. To account for nuclei that appeared to be located eccentrically with respect to the cell cytoplasm, the masks of these nuclei were added to the cell mask upon undergoing a cycle of dilation/erosion aimed at generating a very slim region of surrounding cytoplasm. Myosin outlines were also identified by a global/local-adaptive thresholding procedure but carried out in this case on the G channel alone. Nuclei masks were finally subtracted from the cell and myosin masks.

Once cell, nuclei, cytoplasm, and myosin masks were finalized, the radial distribution of the myosin signal with respect to the centroid of each nucleus was calculated from a radially averaged Difference Patterson map (Patterson map of (nucleus + myosin) – (Patterson map of nucleus + Patterson map of myosin)). The Patterson map [7] of an image is obtained by calculating the fast fourier transform (FFT) of the image, and then back-transforming (inverse FFT, iFFT) removing the phases and using only the squared amplitudes. The map contains all the vectors from every pixel of the image to every other pixel, weighted by the pixel intensity. Thus, subtracting from the Patterson map of the nuclei + myosin (containing nuclei-to-nuclei, nuclei-to-myosin, and myosin-to-myosin vectors) the Patterson map of the nuclei (containing only the nuclei-to-nuclei vectors) and the Patterson map of the myosin (containing only myosin-to-myosin vectors) leaves only the nuclei-to-myosin vectors. Regions of the map with high vector density reflect a strong myosin signal. Radial average of this map provides the distribution of myosin around the nuclei centroids.

Likewise, radial average of a difference Patterson map calculated by subtracting from the Patterson map of all nuclei (containing all vectors between all nuclei) the sum of the Patterson map of an individual nucleus (all vectors inside that nucleus) and the Patterson map of all nuclei except that nucleus provided the distribution of the distances of each nucleus from every other nucleus. Mean nuclei-myosin and nuclei-nuclei density profiles for an entire image were then calculated as the average of the corresponding profiles for each individual nucleus.

### 4. Statistical analysis.

Principal Component Analysis (Fig. 3) and one-way analysis of variance (ANOVA) (Figs. 1,3) were carried out using standard functions of Matlab R2019a Statistics and Machine Learning Toolbox. As to the pdf's of expansion ratios (Fig. 1c), we notice here that the empirical *joint* probability density distribution of nuclei and cell areas before and after expansion is not available because it is not possible to image a sample, expand it, and then re-image it. Instead, the empirical *marginal* distributions (that of unexpanded nuclei/cell sizes and that of expanded nuclei/cell sizes, obtained from different samples) were calculated by kernel density estimation[8] of the observed values of nuclei/cell sizes. Under the assumption of statistical independence, the *joint* pdf was calculated as the outer product of the *marginal* pdf's. Cumulative sum of the probabilities for all observed area ratios (expanded/unexpanded) provided a cumulative distribution function (cdf) of area ratios, which was then converted to the histogram representation of linear expansion ratios (square root of area expansion ratios) shown in Fig. 1c. A normal distribution was then fitted to each histogram by minimizing with respect to mean and standard deviation the norm of the difference between the histogram value and the density of the normal distribution at the center of each bin. A Matlab script streamlining this derivation is provided (see below).

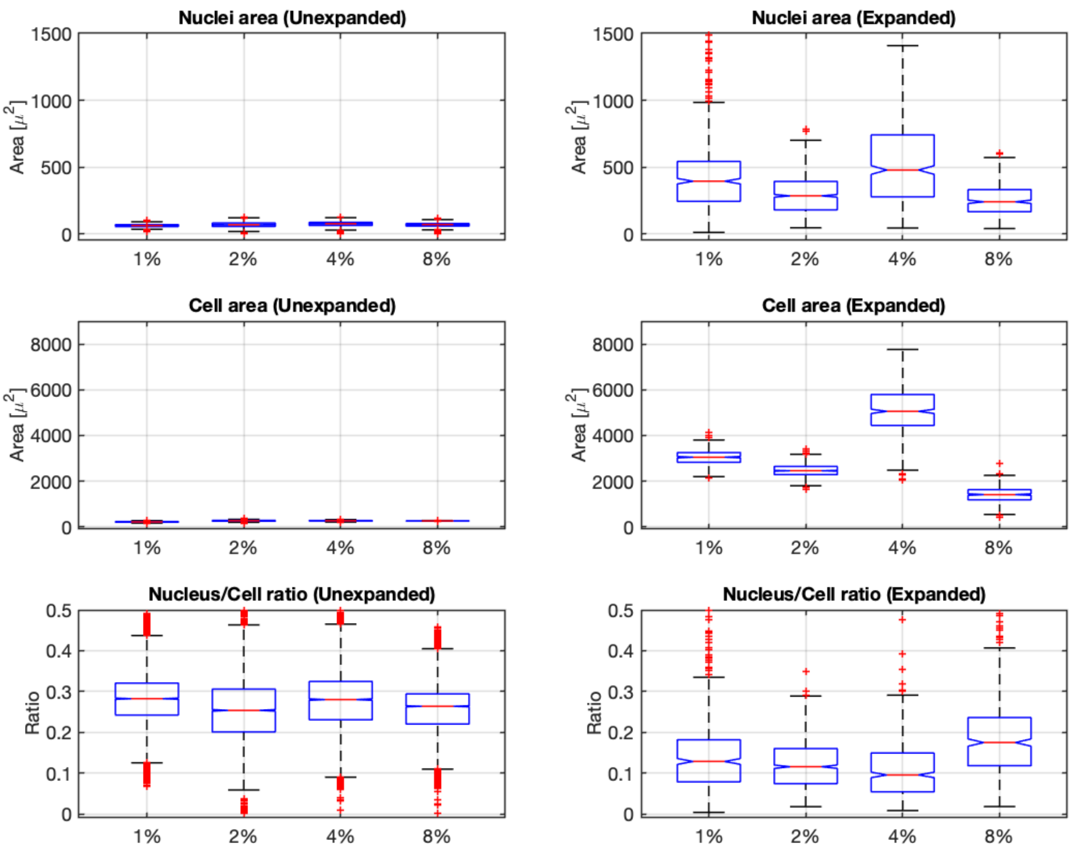


**Figure S1.** Box plots representing one-way analysis of variance (ANOVA) between the distributions of machine learned nuclei and cell features in unexpanded (left column panels) and expanded (right column panels) liver cells at different fixative concentrations. The distribution of nuclei area, cell area, and nucleus to cell area ratio are shown in the top, middle, and bottom row panels, respectively. In each box, the central mark is the median and the edges of the box are the 25th and 75th percentiles (1st and 3rd quantiles). The whiskers extend to the most extreme data points that are not considered outliers. The outliers are plotted individually as red + signs outside the whiskers. In each panel, *p-values* for the joint probability that the distributions at all 4 fixative concentrations share a common mean are all $\ll 10^{-10}$. *p-values* for pairwise one-way AVOVA were $\ll 10^{-10}$ with the exception of the following:

**Control:**

nuclei area 2% *vs* 4% *p-value* = 0.3349

cell area 2% *vs* 3% *p-value* = 0.0038

nuclei/cell ratio 2% *vs* 4% *p-value* = 0.9564

**Expanded:**

nuclei area 2% *vs* 4% *p-value* = 0.0046

nuclei/cell ratio 2% *vs* 4% *p-value* = 0.0751

**5. Neural network recognition of unexpanded and expanded liver cells.**

A neural network capable of accepting as dual inputs DExM microscopy images and numerical data extracted from these images using the Matlab interactive application described above, was build and trained using Keras/Tensorflow [9, 10]. The architecture of the network is shown in Figure S1. Images of 1040 x 1388 pixels were preprocessed to size 260 x 347 pixels prior to 3-fold augmentation by left/right, top/down, and left/right and top/down mirroring combined operations. A total of 45 augmented images belonging to 5 different classes (C1, Control; E1, E2, E4, E8, expanded at 1, 2, 4 and 8% PFA respectively) composed the image training set. The left branch of the network consisted of a convolutional sub-network [4] (CNN) that processed these images, reducing each image to a vector of 128 scalar values. Numerical data associated with the images was in the form of a vector of 805 scalar values containing information on the distributions of nuclei areas, the distribution of nuclei long axis, the distribution of nuclei short axis, the distribution of inter-nuclei distances, the mean cell area, and the mean nucleus to cell area ratio. The right branch of the network consisted of a dense sub-network [11] (DNN) that processed this numerical data and reduced it to a vector of 64 scalar values. Concatenation of the CNN and DNN output vectors resulted in a vector of 192 scalar values, which by application of a *softmax* function resulted in the assignment of each image and the associated numerical data to one of the five classes. Network training was based on the optimization of 122,960 parameters using the *adam* optimizer [12]: random assignment of these parameters at the beginning of the training resulted at the end of 300 *epochs* (with an *epoch* being an optimization cycle encompassing all batches of data in the training set) in a model with 80% class assignment accuracy when tested against 30 images and numerical data never previously seen by the network as part of either training or validation sets. As shown in the *confusion matrix* and the *receiver operating characteristic* [13] (ROC) curves of Figure S3, classes E1 and E8 were not recognized as well as the other three, likely due to their under-representation in the training set.

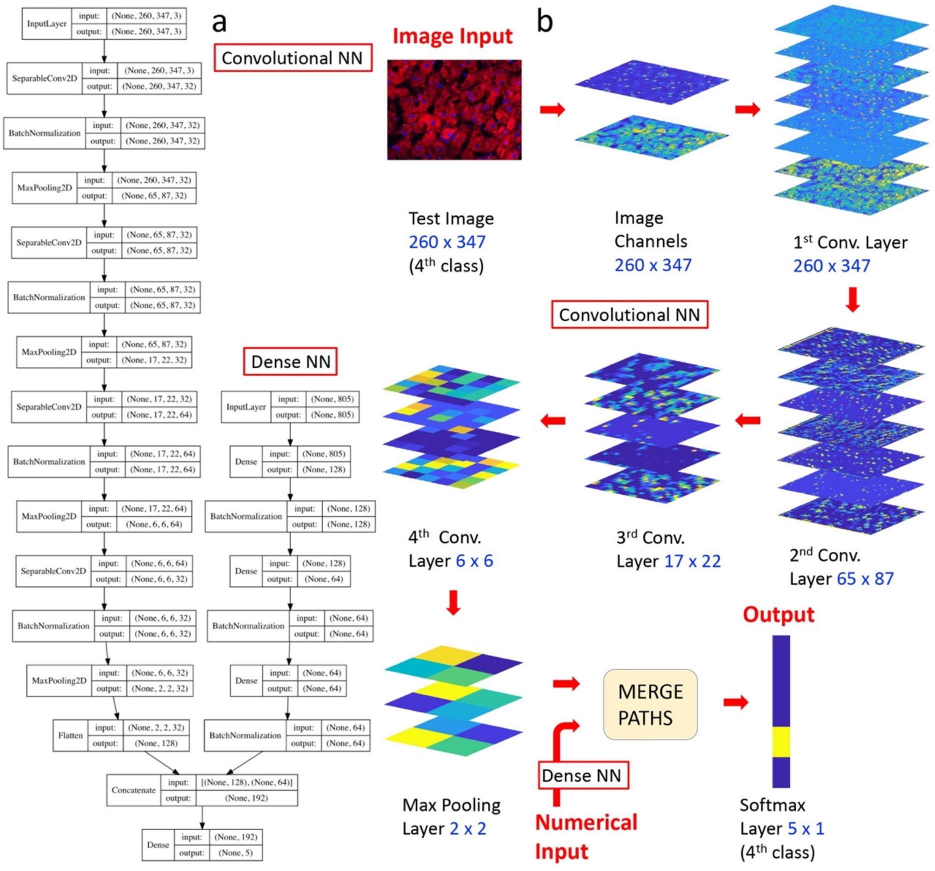


**Figure S2.** Architecture of the mixed-type input NN used to classify images of unexpanded and expanded liver cells. **a.** The left branch of the network consists of a *convolutional* sub-network (CNN) that processes images of 260 x 347 pixels, reducing each image to a vector of 128 scalar values. The right branch of the network consists of a *dense* sub-network (DNN) that processes image associated numerical data in the form of a vector of 805 scalar values and reduces it to a vector of 32 scalar values. Concatenation of the CNN and DNN output vectors results in a vector of 160 scalar values that progresses in a common *dense* path ending with a *softmax* layer for the assignment of each image, and the associated numerical data to one of five classes. **b.** Activations in the CNN branch upon passing a test image of expanded liver cells through the network. Only the activations of some randomly selected channels in each layer are shown. Notice the progressive decrease in the number of pixels (shown in blue font) of the activations, as information is distilled from the initial input image (260 x 347 pixels) down to its class assignment in the softmax layer (5 x 1 pixels).

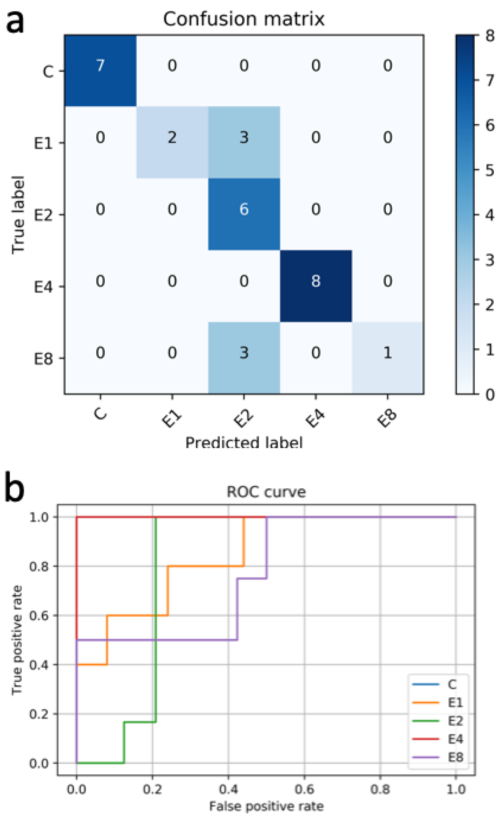


**Figure S3.** **a**: showing the performance of the mixed type (CNN and DNN) classification model. **b**: Receiver operating characteristic curves. Classes: C1, Control; E1, E2, E4, E8, Expanded cells prefixed using 1, 2, 4 and 8 % PFA respectively.

**Video/Audio Tutorial:** http://veloce.med.wayne.edu/~gatti/image-processing.html

**Matlab script to determine the probability distribution of linear expansion ratios:**

```
% Probability distribution of expansion ratios for nuclei and cells. In
% each section of the script the variable Area is used to read a matrix
% with two columns. For example, matrix 'Nuclei_Area_C1_E1' contains in the
% 1st column the area of nuclei in unexpanded cells at 1% fixative (_C1),
% and in the 2nd column the area of nuclei in expanded cells at 1%
% fixative (_E1). The pdf of each feature is calculated using a kernel
% density estimation (function 'ksdensity'). The joint probability
% distribution of these two features is calculated either as the product of
% the marginal distributions (case 'product') or the biased product (case
% 'biased_product').

%% NUCLEI
%%
joint_prob = 'biased_product';

Nuclei_Expansion_distribution = figure;
set(gcf,'Unit','Normalized','Position',[0 0.6 1.0 0.4])

%% 1% Nuclei area

% Marginal distributions of unexpanded and expanded cells
Area = Nuclei_Area_C1_E1;
[fc,xic] = ksdensity(Area(:,1));
[fe,xie] = ksdensity(Area(:,2));
xic_ind = xic >= 0;
xie_ind = xie >= 0;
ind = logical(xic_ind.*xie_ind);
fc = fc(ind);
fe = fe(ind);
xic = xic(ind);
xie = xie(ind);

% figure
% plot(xic,fc,'-r',xie,fe,'-b');grid on
% xlabel('Cell area')
% ylabel('pdf')
% legend('Unexpanded','Expanded')

% Cumulative distribution function of the linear expansions:

%%%%%%%%%%%%%%%%%%%%%%%%%%%%%%%%%%%%%%%%%%%%%%%%%%%%%%%%%%
%%%%%%%%%%%%%%%%
n = 0;
points = zeros(length(xic)^2,2);
for i = 1:length(xic)
    for j = 1:length(xie)
        n = n + 1;
        points(n,:) = [xic(i) xie(j)];
```

```
    end
end

switch joint_prob
    case 'product'
        f = fe'*fc;
    case 'biased_product'
        [f,xi] = ksdensity(Area,points);
end
%%%%%%%%%%%%%%%%%%%%%%%%%%%%%%%%%%%%%%%%%%%%%%%%%%%%%%%%%%%%%%%%%%%%%%%%%%%%%%%%%%%%%%%%%%%%%%%%%%%%%%%%%
%%%%%%%%%%%%%%%%%%%%%%%%%%%%%%%%

% exp_ratios = sqrt(xi(:,2)./xi(:,1));
exp_ratios = sqrt(points(:,2)./points(:,1));
[exp_ratios,sort_order] = sort(exp_ratios);
f = f(sort_order);
f = cumsum(f);
cdf = f/max(f);

% pdf from cdf
subplot(1,4,1)
[n,c] = ecdfhist(cdf,exp_ratios,100);
ecdfhist(cdf,exp_ratios,100); grid on; hold on
h = findobj(gca,'Type','patch');
h.FaceColor = [.8 .8 1];xlabel('Linear expansion ratio')
ylabel('pdf')

% Fit to a normal distribution
fun2 = @(norm_par) norm(n-normpdf(c,norm_par(1),norm_par(2)))
norm_par_0 = [4,1];
norm_par = fminsearch(fun2,norm_par_0);

norm_dist = normpdf(c,norm_par(1),norm_par(2));
plot(c,norm_dist,'r','LineWidth',2)
% grid on
string1 = ['Mean : ' num2str(norm_par(1))];
string2 = ['Std. : ' num2str(norm_par(2))];
min_length = min([length(string1) length(string2)]);
string1 = string1(1:min_length);
string2 = string2(1:min_length);
annotation('textbox', [0.2, 0.7, 0.3, 0.1], 'String', [string1; string2], ...
    'FitBoxToText','on','Color','red','FontSize',14,'EdgeColor','b')

xlabel('Linear expansion ratio')
ylabel('pdf')


%% 2% Nuclei area

% Marginal distributions of unexpanded and expanded cells
```

```
Area = Nuclei_Area_C2_E2;
[fc,xic] = ksdensity(Area(:,1));
[fe,xie] = ksdensity(Area(:,2));
xic_ind = xic >= 0;
xie_ind = xie >= 0;
ind = logical(xic_ind.*xie_ind);
fc = fc(ind);
fe = fe(ind);
xic = xic(ind);
xie = xie(ind);

% figure
% plot(xic,fc,'-r',xie,fe,'-b');grid on
% xlabel('Cell area')
% ylabel('pdf')
% legend('Unexpanded','Expanded')

% Cumulative distribution function of the linear expansions

%%%%%%%%%%%%%%%%%%%%%%%%%%%%%%%%%%%%%%%%%%%%%%%%%%%%%%%%%%%%%%%%%%%
%%%%%%%%%%%%%%%%%%
n = 0;
points = zeros(length(xic)^2,2);
for i = 1:length(xic)
    for j = 1:length(xie)
        n = n + 1;
        points(n,:) = [xic(i) xie(j)];
    end
end

switch joint_prob
    case 'product'
        f = fe'*fc;
    case 'biased_product'
        [f,xi] = ksdensity(Area,points);
end
%%%%%%%%%%%%%%%%%%%%%%%%%%%%%%%%%%%%%%%%%%%%%%%%%%%%%%%%%%%%%%%%%%%
%%%%%%%%%%%%%%%%%%

% exp_ratios = sqrt(xi(:,2)./xi(:,1));
exp_ratios = sqrt(points(:,2)./points(:,1));
[exp_ratios,sort_order] = sort(exp_ratios);
f = f(sort_order);
f = cumsum(f);
cdf = f/max(f);

% pdf from cdf
subplot(1,4,2)
[n,c] = ecdfhist(cdf,exp_ratios,100);
ecdfhist(cdf,exp_ratios,100); grid on; hold on
```

```
h = findobj(gca,'Type','patch');
h.FaceColor = [.8 1 .8];xlabel('Linear expansion ratio')
ylabel('pdf')

% Fit to a normal distribution
fun2 = @(norm_par) norm(n-normpdf(c,norm_par(1),norm_par(2)))
norm_par_0 = [4,1];
norm_par = fminsearch(fun2,norm_par_0);

norm_dist = normpdf(c,norm_par(1),norm_par(2));
plot(c,norm_dist,'r','LineWidth',2)
% grid on
string1 = ['Mean : ' num2str(norm_par(1))];
string2 = ['Std. : ' num2str(norm_par(2))];
min_length = min([length(string1) length(string2)]);
string1 = string1(1:min_length);
string2 = string2(1:min_length);
annotation('textbox', [0.415, 0.7, 0.3, 0.1], 'String', [string1; string2], ...
    'FitBoxToText','on','Color','red','FontSize',14,'EdgeColor','b')

xlabel('Linear expansion ratio')
ylabel('pdf')

%% 4% Nuclei area

% Marginal distributions of unexpanded and expanded cells
Area = Nuclei_Area_C4_E4;
[fc,xic] = ksdensity(Area(:,1));
[fe,xie] = ksdensity(Area(:,2));
xic_ind = xic >= 0;
xie_ind = xie >= 0;
ind = logical(xic_ind.*xie_ind);
fc = fc(ind);
fe = fe(ind);
xic = xic(ind);
xie = xie(ind);

% figure
% plot(xic,fc,'-r',xie,fe,'-b');grid on
% xlabel('Cell area')
% ylabel('pdf')
% legend('Unexpanded','Expanded')

% Cumulative distribution function of the linear expansions

%%%%%%%%%%%%%%%%%%%%%%%%%%%%%%%%%%%%%%%%%%%%%%%%%%%%%%%%%%%%%%%%%%%%%%%%%%%%%%%%%%%
%%%%%%%%%%%%%%%%%%%%%
n = 0;
points = zeros(length(xic)^2,2);
for i = 1:length(xic)
```

```
    for j = 1:length(xie)
        n = n + 1;
        points(n,:) = [xic(i) xie(j)];
    end
end

switch joint_prob
    case 'product'
        f = fe'*fc;
    case 'biased_product'
        [f,xi] = ksdensity(Area,points);
end
%%%%%%%%%%%%%%%%%%%%%%%%%%%%%%%%%%%%%%%%%%%%%%%%
%%%%%%%%%%%%%%%%%

% exp_ratios = sqrt(xi(:,2)./xi(:,1));
exp_ratios = sqrt(points(:,2)./points(:,1));
[exp_ratios,sort_order] = sort(exp_ratios);
f = f(sort_order);
f = cumsum(f);
cdf = f/max(f);

% pdf from cdf
subplot(1,4,3)
[n,c] = ecdfhist(cdf,exp_ratios,100);
ecdfhist(cdf,exp_ratios,100); grid on; hold on
h = findobj(gca,'Type','patch');
h.FaceColor = [1 .8 .8];xlabel('Linear expansion ratio')
ylabel('pdf')

% Fit to a normal distribution
fun2 = @(norm_par) norm(n-normpdf(c,norm_par(1),norm_par(2)))
norm_par_0 = [4,1];
norm_par = fminsearch(fun2,norm_par_0);

norm_dist = normpdf(c,norm_par(1),norm_par(2));
plot(c,norm_dist,'r','LineWidth',2)
% grid on
string1 = ['Mean : ' num2str(norm_par(1))];
string2 = ['Std. : ' num2str(norm_par(2))];
min_length = min([length(string1) length(string2)]);
string1 = string1(1:min_length);
string2 = string2(1:min_length);
annotation('textbox', [0.6229, 0.7, 0.3, 0.1], 'String', [string1; string2], ...
    'FitBoxToText','on','Color','red','FontSize',14,'EdgeColor','b')

xlabel('Linear expansion ratio')
ylabel('pdf')

%% 8% Nuclei area
```

```
% Marginal distributions of unexpanded and expanded cells
Area = Nuclei_Area_C8_E8;
[fc,xic] = ksdensity(Area(:,1));
[fe,xie] = ksdensity(Area(:,2));
xic_ind = xic >= 0;
xie_ind = xie >= 0;
ind = logical(xic_ind.*xie_ind);
fc = fc(ind);
fe = fe(ind);
xic = xic(ind);
xie = xie(ind);

% figure
% plot(xic,fc,'-r',xie,fe,'-b');grid on
% xlabel('Cell area')
% ylabel('pdf')
% legend('Unexpanded','Expanded')

% Cumulative distribution function of the linear expansions

%%%%%%%%%%%%%%%%%%%%%%%%%%%%%%%%%%%%%%%%%%
%%%%%%%%%%%%%%
n = 0;
points = zeros(length(xic)^2,2);
for i = 1:length(xic)
    for j = 1:length(xie)
        n = n + 1;
        points(n,:) = [xic(i) xie(j)];
    end
end

switch joint_prob
    case 'product'
        f = fe'*fc;
    case 'biased_product'
        [f,xi] = ksdensity(Area,points);
end
%%%%%%%%%%%%%%%%%%%%%%%%%%%%%%%%%%%%%%%%%%
%%%%%%%%%%%%%%

% exp_ratios = sqrt(xi(:,2)./xi(:,1));
exp_ratios = sqrt(points(:,2)./points(:,1));
[exp_ratios,sort_order] = sort(exp_ratios);
f = f(sort_order);
f = cumsum(f);
cdf = f/max(f);

% pdf from cdf
subplot(1,4,4)
```

```
[n,c] = ecdfhist(cdf,exp_ratios,100);
ecdfhist(cdf,exp_ratios,100); grid on; hold on
h = findobj(gca,'Type','patch');
h.FaceColor = [.8 .8 .8];xlabel('Linear expansion ratio')
ylabel('pdf')

% Fit to a normal distribution
fun2 = @(norm_par) norm(n-normpdf(c,norm_par(1),norm_par(2)))
norm_par_0 = [4,1];
norm_par = fminsearch(fun2,norm_par_0);

norm_dist = normpdf(c,norm_par(1),norm_par(2));
plot(c,norm_dist,'r','LineWidth',2)
% grid on
string1 = ['Mean : ' num2str(norm_par(1))];
string2 = ['Std. : ' num2str(norm_par(2))];
min_length = min([length(string1) length(string2)]);
string1 = string1(1:min_length);
string2 = string2(1:min_length);
annotation('textbox', [0.8333, 0.7, 0.3, 0.1], 'String', [string1; string2], ...
   'FitBoxToText','on','Color','red','FontSize',14,'EdgeColor','b')

xlabel('Linear expansion ratio')
ylabel('pdf')

%% CELLS
joint_prob = 'product';

Cell_Expansion_distribution = figure;
set(gcf,'Unit','Normalized','Position',[0 0.6 1.0 0.4])

%% 1% cell area

% Marginal distributions of unexpanded and expanded cells
Area = Cell_Area_C1_E1;
[fc,xic] = ksdensity(Area(:,1));
[fe,xie] = ksdensity(Area(:,2));
xic_ind = xic >= 0;
xie_ind = xie >= 0;
ind = logical(xic_ind.*xie_ind);
fc = fc(ind);
fe = fe(ind);
xic = xic(ind);
xie = xie(ind);

% figure
% plot(xic,fc,'-r',xie,fe,'-b');grid on
% xlabel('Cell area')
% ylabel('pdf')
% legend('Unexpanded','Expanded')
```

```
% Cumulative distribution function of the linear expansions:

%%%%%%%%%%%%%%%%%%%%%%%%%%%%%%%%%%%%%%%%%%%%%%%%%%%%%
%%%%%%%%%%%%%%%%%%%%%%%
n = 0;
points = zeros(length(xic)^2,2);
for i = 1:length(xic)
   for j = 1:length(xie)
      n = n + 1;
      points(n,:) = [xic(i) xie(j)];
   end
end

switch joint_prob
   case 'product'
      f = fe'*fc;
   case 'biased_product'
      [f,xi] = ksdensity(Area,points);
end
%%%%%%%%%%%%%%%%%%%%%%%%%%%%%%%%%%%%%%%%%%%%%%%%%%%%%
%%%%%%%%%%%%%%%%%%%%%%%

% exp_ratios = sqrt(xi(:,2)./xi(:,1));
exp_ratios = sqrt(points(:,2)./points(:,1));
[exp_ratios,sort_order] = sort(exp_ratios);
f = f(sort_order);
f = cumsum(f);
cdf = f/max(f);

% pdf from cdf
subplot(1,4,1)
[n,c] = ecdfhist(cdf,exp_ratios,100);
ecdfhist(cdf,exp_ratios,100); grid on; hold on
h = findobj(gca,'Type','patch');
h.FaceColor = [.8 .8 1];xlabel('Linear expansion ratio')
ylabel('pdf')

% Fit to a normal distribution
fun2 = @(norm_par) norm(n-normpdf(c,norm_par(1),norm_par(2)))
norm_par_0 = [4,1];
norm_par = fminsearch(fun2,norm_par_0);

norm_dist = normpdf(c,norm_par(1),norm_par(2));
plot(c,norm_dist,'r','LineWidth',2)
% grid on
string1 = ['Mean : ' num2str(norm_par(1))];
string2 = ['Std. : ' num2str(norm_par(2))];
min_length = min([length(string1) length(string2)]);
string1 = string1(1:min_length);
```

```
string2 = string2(1:min_length);
annotation('textbox', [0.2, 0.7, 0.3, 0.1], 'String', [string1; string2], ...
   'FitBoxToText','on','Color','red','FontSize',14,'EdgeColor','b')

xlabel('Linear expansion ratio')
ylabel('pdf')


%% 2% cell area

% Marginal distributions of unexpanded and expanded cells
Area = Cell_Area_C2_E2;
[fc,xic] = ksdensity(Area(:,1));
[fe,xie] = ksdensity(Area(:,2));
xic_ind = xic >= 0;
xie_ind = xie >= 0;
ind = logical(xic_ind.*xie_ind);
fc = fc(ind);
fe = fe(ind);
xic = xic(ind);
xie = xie(ind);

% figure
% plot(xic,fc,'-r',xie,fe,'-b');grid on
% xlabel('Cell area')
% ylabel('pdf')
% legend('Unexpanded','Expanded')

% Cumulative distribution function of the linear expansions

%%%%%%%%%%%%%%%%%%%%%%%%%%%%%%%%%%%%%%%%%%%%
%%%%%%%%%%%%%%%
n = 0;
points = zeros(length(xic)^2,2);
for i = 1:length(xic)
   for j = 1:length(xie)
      n = n + 1;
      points(n,:) = [xic(i) xie(j)];
   end
end

switch joint_prob
   case 'product'
      f = fe'*fc;
   case 'biased_product'
      [f,xi] = ksdensity(Area,points);
end
%%%%%%%%%%%%%%%%%%%%%%%%%%%%%%%%%%%%%%%%%%%%
%%%%%%%%%%%%%%%
```

```
% exp_ratios = sqrt(xi(:,2)./xi(:,1));
exp_ratios = sqrt(points(:,2)./points(:,1));
[exp_ratios,sort_order] = sort(exp_ratios);
f = f(sort_order);
f = cumsum(f);
cdf = f/max(f);

% pdf from cdf
subplot(1,4,2)
[n,c] = ecdfhist(cdf,exp_ratios,100);
ecdfhist(cdf,exp_ratios,100); grid on; hold on
h = findobj(gca,'Type','patch');
h.FaceColor = [.8 1 .8];xlabel('Linear expansion ratio')
ylabel('pdf')

% Fit to a normal distribution
fun2 = @(norm_par) norm(n-normpdf(c,norm_par(1),norm_par(2)))
norm_par_0 = [4,1];
norm_par = fminsearch(fun2,norm_par_0);

norm_dist = normpdf(c,norm_par(1),norm_par(2));
plot(c,norm_dist,'r','LineWidth',2)
% grid on
string1 = ['Mean : ' num2str(norm_par(1))];
string2 = ['Std. : ' num2str(norm_par(2))];
min_length = min([length(string1) length(string2)]);
string1 = string1(1:min_length);
string2 = string2(1:min_length);
annotation('textbox', [0.415, 0.7, 0.3, 0.1], 'String', [string1; string2], ...
    'FitBoxToText','on','Color','red','FontSize',14,'EdgeColor','b')

xlabel('Linear expansion ratio')
ylabel('pdf')

%% 4% cell area

% Marginal distributions of unexpanded and expanded cells
Area = Cell_Area_C4_E4;
[fc,xic] = ksdensity(Area(:,1));
[fe,xie] = ksdensity(Area(:,2));
xic_ind = xic >= 0;
xie_ind = xie >= 0;
ind = logical(xic_ind.*xie_ind);
fc = fc(ind);
fe = fe(ind);
xic = xic(ind);
xie = xie(ind);

% figure
% plot(xic,fc,'-r',xie,fe,'-b');grid on
```

```
% xlabel('Cell area')
% ylabel('pdf')
% legend('Unexpanded','Expanded')

% Cumulative distribution function of the linear expansions

%%%%%%%%%%%%%%%%%%%%%%%%%%%%%%%%%%%%%%%%%%%%%%%%%%%%%
%%%%%%%%%%%%%%%%
n = 0;
points = zeros(length(xic)^2,2);
for i = 1:length(xic)
   for j = 1:length(xie)
      n = n + 1;
      points(n,:) = [xic(i) xie(j)];
   end
end

switch joint_prob
   case 'product'
      f = fe'*fc;
   case 'biased_product'
      [f,xi] = ksdensity(Area,points);
end
%%%%%%%%%%%%%%%%%%%%%%%%%%%%%%%%%%%%%%%%%%%%%%%%%%%%%
%%%%%%%%%%%%%%%%

% exp_ratios = sqrt(xi(:,2)./xi(:,1));
exp_ratios = sqrt(points(:,2)./points(:,1));
[exp_ratios,sort_order] = sort(exp_ratios);
f = f(sort_order);
f = cumsum(f);
cdf = f/max(f);

% pdf from cdf
subplot(1,4,3)
[n,c] = ecdfhist(cdf,exp_ratios,100);
ecdfhist(cdf,exp_ratios,100); grid on; hold on
h = findobj(gca,'Type','patch');
h.FaceColor = [1 .8 .8];xlabel('Linear expansion ratio')
ylabel('pdf')

% Fit to a normal distribution
fun2 = @(norm_par) norm(n-normpdf(c,norm_par(1),norm_par(2)))
norm_par_0 = [4,1];
norm_par = fminsearch(fun2,norm_par_0);

norm_dist = normpdf(c,norm_par(1),norm_par(2));
plot(c,norm_dist,'r','LineWidth',2)
% grid on
string1 = ['Mean : ' num2str(norm_par(1))];
```

```
string2 = ['Std. : ' num2str(norm_par(2))];
min_length = min([length(string1) length(string2)]);
string1 = string1(1:min_length);
string2 = string2(1:min_length);
annotation('textbox', [0.6229, 0.7, 0.3, 0.1], 'String', [string1; string2], ...
   'FitBoxToText','on','Color','red','FontSize',14,'EdgeColor','b')

xlabel('Linear expansion ratio')
ylabel('pdf')

%% 8% cell area

% Marginal distributions of unexpanded and expanded cells
Area = Cell_Area_C8_E8;
[fc,xic] = ksdensity(Area(:,1));
[fe,xie] = ksdensity(Area(:,2));
xic_ind = xic >= 0;
xie_ind = xie >= 0;
ind = logical(xic_ind.*xie_ind);
fc = fc(ind);
fe = fe(ind);
xic = xic(ind);
xie = xie(ind);

% figure
% plot(xic,fc,'-r',xie,fe,'-b');grid on
% xlabel('Cell area')
% ylabel('pdf')
% legend('Unexpanded','Expanded')

% Cumulative distribution function of the linear expansions

%%%%%%%%%%%%%%%%%%%%%%%%%%%%%%%%%%%%%%%%%%%%%
%%%%%%%%%%%%%%%
n = 0;
points = zeros(length(xic)^2,2);
for i = 1:length(xic)
   for j = 1:length(xie)
      n = n + 1;
      points(n,:) = [xic(i) xie(j)];
   end
end

switch joint_prob
   case 'product'
      f = fe'*fc;
   case 'biased_product'
      [f,xi] = ksdensity(Area,points);
end
```

```
%%%%%%%%%%%%%%%%%%%%%%%%%%%%%%%%%%%%%%%%%%%%%%%%%%%%%%%%%%%%%%%%%%%%%%%%%%%%%

% exp_ratios = sqrt(xi(:,2)./xi(:,1));
exp_ratios = sqrt(points(:,2)./points(:,1));
[exp_ratios,sort_order] = sort(exp_ratios);
f = f(sort_order);
f = cumsum(f);
cdf = f/max(f);

% pdf from cdf
subplot(1,4,4)
[n,c] = ecdfhist(cdf,exp_ratios,100);
ecdfhist(cdf,exp_ratios,100); grid on; hold on
h = findobj(gca,'Type','patch');
h.FaceColor = [.8 .8 .8];xlabel('Linear expansion ratio')
ylabel('pdf')

% Fit to a normal distribution
fun2 = @(norm_par) norm(n-normpdf(c,norm_par(1),norm_par(2)))
norm_par_0 = [4,1];
norm_par = fminsearch(fun2,norm_par_0);

norm_dist = normpdf(c,norm_par(1),norm_par(2));
plot(c,norm_dist,'r','LineWidth',2)
% grid on
string1 = ['Mean : ' num2str(norm_par(1))];
string2 = ['Std. : ' num2str(norm_par(2))];
min_length = min([length(string1) length(string2)]);
string1 = string1(1:min_length);
string2 = string2(1:min_length);
annotation('textbox', [0.8333, 0.7, 0.3, 0.1], 'String', [string1; string2], ...
    'FitBoxToText','on','Color','red','FontSize',14,'EdgeColor','b')

xlabel('Linear expansion ratio')
ylabel('pdf')
```
